# Self-reported sleep duration and daytime napping are associated with renal hyperfiltration and microalbuminuria in apparently healthy Chinese population

**DOI:** 10.1101/585190

**Authors:** Yingnan Ye, Linxi Zhang, Wenhua Yan, Anping Wang, Weiqing Wang, Zhengnan Gao, Xulei Tang, Li Yan, Qin Wan, Zuojie Luo, Guijun Qin, Lulu Chen, Shiqing Wang, Yuxia Wang, Yiming Mu

**Affiliations:** Chinese PLA General Hospital, Beijing, China; Department of Medicine, Nankai University, Tianjin, China; Shanghai jiaotong University Affiliated Ruijin Hospital, ShangHai, China; Center Hospital of Dalian, Dalian, Liaoning, China; Lanzhou University first Hospital, Lanzhou, Gansu, China; Zhongshan University Sun yat-sen memorial Hospital, Guangzhou, Guangdong, China; Southwest Medical University Affiliated Hospital, Luzhou, Sichuan, China; Guangxi Medical University first affiliated Hospital, Nanning, Guangxi, China; Zhengzhou University first affiliated Hospital, Zhengzhou, Henan, China; Wuhan Union Hospital, Wuhan, Hubei, China

**Keywords:** Sleep duration, daytime napping, albuminuria, hyperfiltration, renal health

## Abstract

**Background:** Sleep duration affects health in various way. The objective of this study was to investigate the relationship between sleep duration, daytime napping and kidney function in a middle-aged apparently healthy Chinese population.

**Methods:** According to self-reported total sleep and daytime napping duration, 33,850 participants aged 38 to 90 years old from 8 regional centers were divided into subgroups. Height, weight, waistline, hipline, blood pressure, biochemical index, FBG, PBG, HbA1c, creatinine and urinary albumin-creatinine ratio (UACR) were measured and recorded in each subject. Microalbuminuria was defined as UACR>=30 mg/g, CKD was defined as eGFR<60 ml/min and hyperfiltration was defined as eGFR>=135 ml/min. Multiple logistic regressions were applied to investigate associations between sleep and kidney function.

**Results:** Compared to participants with [7-8]h/day sleep, ORs of >9 h/day, (8, 9]h/day and <6h/day sleep for microalbuminuria were 1.317 (1.200-1.446, p<0.001), 1.215 (1.123-1.315, p<0.001) and 1.218 (0.967-1.534, p=0.094). eGFR levels were U-shaped associated with sleep duration among subjects with >=90ml/min eGFR, and N-shaped associated with sleep duration among subjects with <90ml/min eGFR. OR of >9h/day sleep for hyperfiltration was 1.400 (1.123-1.745, p=0.003) among eGFR>=90 ml/min participants. Daytime napping had a negative effect on renal health. Compared to participants did not have napping habit, the ORs of (0, 1]h/day, (1, 1.5]h/day and >1.5h/day daytime napping for microalbuminuria were 1.477 (1.370-1.591, p<0.001), 1.217 (1.056, 1.403, p=0.007) and 1.447 (1.242, 1.687, p<0.001).

**Conclusions:** Total sleep duration are U-shaped associated with renal health outcomes. Daytime napping had a negative effect on renal health.

## Introduction

In the past decades, accumulating evidences indicated chronic sleep disorders represent a risk factor affecting metabolic health. Inappropriate sleep duration had been proved to be associated with many adverse health outcomes, such as diabetes^1, 2^, obesity^3^, hypertension^4, 5^, osteoporosis^6^, cardiovascular disease^7, 8^, stroke^9^ and total mortality^10^. Recently, a series of studies suggested extreme sleep duration may contribute to the decline of kidney function, which is closely tied to the vascular system, also an important, independent risk factor for cardiovascular disease. Both extremely short^11, 12^ and long^13^ sleep duration and poor sleep quality^14, 26^ were reported to be related to higher urine albumin-to-creatinine ratio(UACR), which is a sensitive indicator for microalbuminuria or early stage of kidney damage, among US and Japanese population. In addition, short sleep duration was associated with higher odds of inadequate hydration^15^. On the other hand, whether the effect of sleep duration on glomerular filtration rate(GFR) is positive or negative is debatable. Several studies have demonstrated an increasing risk of CKD16,17 or lower GFR12,18,19,20,21 in short sleepers, but a few studies have shown no correlation between sleep duration and CKD22-25. Conversely, there were both cross-sectional or cohort studies which have reported inappropriate sleep duration contributes to glomerular hyperfiltration^13, 26–28^. This difference in outcomes can be attributed not only to differences in race, age, social work stress, health and economic status of the participants, but also to the fact that in the progression of CKD, healthy individuals tend to have glomerular hyperfiltration initially, followed by increased risk for renal injury, leading to a decrease in filtration rate and accelerating development of CKD. Such a pathophysiological progression occurs in the context of type 1 diabetes and hypertension, as well as increasing stages of pre-diabetes and pre-hypertension^29–31^. Therefore, to provide further evidence of the contribution of sleep duration in the progression of renal function decline, a multi-center studies with sufficient participants with diverse health conditions will be required. Therefore, we conducted this study to determine whether the relationship between UACR and sleep duration exists among Chinese provinces and cities, and to verify whether different health conditions have an interactive effect on this relationship.

## Methods

### Study subjects

A total of 33,850 participants from 8 regional centers include in the REACTION (Risk Evaluation of cAncers in Chinese diabeTic Individuals a LONgitudinal) study, in which are Dalian, Guangzhou, Zhengzhou, Lanzhou, Luzhou, Wuhan, Guangxi, and Shanghai. Excluded participants with primary kidney diseases, daily ACEI/ARB medicine use, and those with a fallacious self-reported sleep duration (<4 h or >12 h).

### Questionnaire

A standardized questionnaire was used to collect basic information including medical history, physical exercise, and smoking and drinking habits. Self-reported sleep duration and daytime napping time were ascertained by following questions: (1) how many hours of sleep do you usually get at night, (2) how many minutes do you usually nap at noon. All investigators were previously trained. In the analysis, participants were divided into four groups (<6, >=6&<7, >=7&<=8, >8%<=9, >9 h/d) on the basis of their sleep duration for 7 to 8 hours of sleep is generally considered as the most appropriate sleep duration.

### Physical examination

Height, weight, waistline, and hipline of the subjects were measured and recorded. Participants were asked to take off shoes, hats and coats before measurements. Waistline was measured at the horizontal level of the midpoint of the ligature between anterior superior spine and the inferior margin of the 12th rib. Hipline was defined as the horizontal length of the most protruding part of hip. All data were recorded to within one decimal place.

### Urinary albumin-creatinine ratio (UACR) measurement and data processing

Urine samples were collected in the morning for the UACR measurement. According to the quartile division, among which centre the subjects belonged in the logistic regression, the UACR data were divided into the under 25% group, the 25%-50% group, the 50%-75% group, and the over 75% group. Higher UACR level was defined as subjects belonged to the over 75% group. Microalbuminuria was defined as UACR>=30 mg/g.

### Estimated glomerular filtration rate calculation

MDRD formula was used to calculate eGFR.
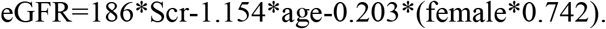

CKD was defined as eGFR<60 ml/min. Hyperfiltration was defined as eGFR>=135 ml/min.

### Blood pressure measurement

Blood pressure was measured 3 times with 1 min intervals after subjects were seated for 5 min. The average of 3 values was used for the analysis. Hypertension was defined as the average systolic blood pressure >=140 mmHg or diastolic blood pressure >=90 mmHg, or had a definite medical history of hypertension.

### Blood biochemical index, glucose, HbA1c and insulin measurement

Blood samples were drawn in the morning after subjects had fasted 8 hours the previous night. Participants without a history of diabetes underwent a 75-g oral glucose tolerance test, while those with diabetes underwent a 100g oral steam bread tolerance test and their venous blood samples were drawn at 0 and 120 minutes. Biochemical index included triglycerides (TG), cholesterol (TC), low density lipoprotein (LDL), high density lipoprotein (HDL), creatinine (Scr), urea nitrogen (BUN), liver function index (ALT, AST, GGT), fasting blood glucose (FBG), postprandial blood glucose (PBG), glycosylated haemoglobin (HbA1c), and fasting blood insulin and postprandial blood insulin, which were measured by glucose oxidase-peroxidase method. Diabetes mellitus (DM) was defined as FBG>=7.0 mmol/L or PBG>=11.1mmol/L, or had a definite medical history of diabetes. Impaired fasting glucose was defined as FBG>=6.1 mmol/L but without DM. Impaired glucose tolerance was defined as PBG>=7.8 mmol/L but without DM.

### Statistical analysis

Statistical analysis was performed using SPSS software, version 19.0 (Chicago, IL). All continuous variables with normal distriution are presented as the mean values and standard deviation (SD). All continuous variables with skewness distribution are presented as media and 25,75 percentile. All enumeration data presented as propotion. The differences in the mean values or proportions of the characteristics of the studied subjects sectional association between sleep duration and UACR, FPG, PPG, AST, GGT, ALT, HbA1c and TG values, and the latter were transformed using the natural log in the analyses due to their skewed distribution.

## Results

A total of 33,850 participants, including 11,198 males and 22,652 females were included in the analysis. The mean age of the total population was 58.1±9.3 years; self-reported total sleep duration was 8.1±1.2 h, and daytime napping duration was 0.3±0.6 h; median of UACR was 9.4 mg/g; mean eGFR was 93.9±19.1 ml/min.

General characteristics of the participants in this study according to different categories of total sleep duration are shown in Table 1. Compared with those who slept 7-8 h per day, participants who slept shorter or longer were more likely to be older and to have higher TG, PBG and UACR value, as well as higher proportion of diabetes, hypertension, hyperfiltration, micoalbuminuria and high UACR level. Those with sleep duration longer than 9 h were more likely to be obesity and to have lower eGFR level and less physic exercise, however, their CHOL and LDL levels were lower compared to the reference. Reverse results were observed in the short sleep duration groups. Furthermore, no significant difference in blood pressure was observed across all sleep categories. Although there were statistical differences in drinking and smoking habits across the groups, no significant increase or decrease or U-shaped relationship between sleep duration and drinking or smoking was observed.

**Table 1.** Baseline information according to self-reported total sleep duration.

| Sleep duration,h | ALL n=33850 | (0- 6) n=651 | [6-7) n=2854 | [7--8] n=16492 | (8--9] n=8902 | >9 n=4951 | P value |
| --- | --- | --- | --- | --- | --- | --- | --- |
| Age, years | 58.1±9.3 | 58.7±8.8 | 57.7±8.8 | 57.5±8.8 | 58.5±9.6 | 59.8±10.3 | <0.001 |
| Male, % | 33.1(11198) | 31.5(205) | 31.7(904) | 32.1(5291) | 33.7(2997) | 36.4(1801) | <0.001 |
| Waist circumference,cm | 86.2±10.0 | 87.6±11.2 | 86.4±10.1 | 86.0±9.9 | 86.3±10.0 | 86.4±10.2 | <0.001 |
| Hip circumference,cm | 97.0±7.8 | 97.7±8.6 | 97.3±7.7 | 97.0±7.8 | 97.0±7.8 | 96.6±7.9 | <0.001 |
| BMI, kg/m2 | 24.7±3.7 | 25.1±4.3 | 24.9±3.6 | 24.7±3.7 | 24.6±3.6 | 24.5±3.7 | <0.001 |
| Tumor,% | 3.0(992) | 3.7(24) | 3.3(93) | 2.9(485) | 2.8(252) | 2.8(138) | 0.515 |
| Diabetes,% | 22.8(7712) | 24.7(161) | 21.8(622) | 21.0(3461) | 23.4(2082) | 28.0(1386) | <0.001 |
| Smoking habits |  |  |  |  |  |  | <0.001 |
| Regular smoker,% | 12.6(4219) | 13.2(85) | 13.2(372) | 12.1(1983) | 12.2(1075) | 14.3(704) |  |
| Sometimes smoker,% | 2.6(878) | 1.4(9) | 2.4(68) | 2.5(407) | 2.9(257) | 2.8(137) |  |
| Never smoker,% | 84.8(28470) | 85.4(551) | 83.5(2384) | 85.4(13975) | 84.9(7493) | 82.9(4067) |  |
| Drinking habits |  |  |  |  |  |  | <0.001 |
| Regular drinker,% | 7.0(2349) | 9.6(62) | 7.7(219) | 6.6(1080) | 7.1(626) | 7.4(362) |  |
| Sometimes drinker,% | 19.1(6406) | 17.0(110) | 19.8(560) | 19.9(3262) | 17.9(1580) | 18.2(894) |  |
| Never drinker,% | 73.9(24827) | 73.4(475) | 72.4(2048) | 73.5(12034) | 75.0(6620) | 74.4(3650) |  |
| Regular exercise,% | 12.1(4097) | 14.0(91) | 12.9(368) | 12.9(2133) | 11.3(1008) | 9.9(497) | <0.001 |
| Exercise intensity |  |  |  |  |  |  | <0.001 |
| Mild exercise,% | 9.1(3081) | 9.8(64) | 9.7(278) | 10.2(1675) | 8.4(750) | 6.3(314) |  |
| Moderate exercise,% | 2.6(875) | 3.4(22) | 2.8(80) | 2.4(403) | 2.5(224) | 2.9(146) |  |
| Severe exercise,% | 0.4(141) | 0.8(5) | 0.4(10) | 0.3(55) | 0.4(34) | 0.7(37) |  |
| Obesity(BMI>=28),% | 15.1(5127) | 18.9(123) | 17.4(2854) | 15.2(2510) | 14.5(1293) | 14.2(703) | <0.001 |
| HbA1c,% | 5.9(5.6,6.2) | 5.9(5.6,6.3) | 5.9(5.6,6.2) | 5.9(5.6,6.2) | 5.9(5.6,6.3) | 5.9(5.6,6.4) | <0.001 |
| SBP,mmHg | 131.8±20.4 | 132.5±22.0 | 132.1±20.2 | 131.9±20.3 | 131.6±20.4 | 131.9±20.8 | 0.695 |
| DBP,mmHg | 77.6±10.9 | 77.4±11.1 | 77.6±11.0 | 77.6±10.9 | 77.5±10.8 | 77.6±10.9 | 0.795 |
| Hypertension,% | 40.5(13708) | 43.6(284) | 39.7(1133) | 39.6(6523) | 40.9(3643) | 42.9(2125) | <0.001 |
| Menopause(Female),% | 74.3(14270) | 79.3(280) | 74.5(1178) | 88.4(14580) | 74.7(3830) | 75.8(2100) | 0.019 |
| CreaC,mmol/L | 65.8(59.5,73.7) | 65.3(59.1,72.6) | 65.2(59.1,73.2) | 65.4(59.3,73.1) | 65.9(59.5,74.2) | 67.0(60.3,75.8) | <0.001 |
| Triglycerides,mmol/L | 1.36(0.97,1.97) | 1.36(0.98,1.96) | 1.32(0.92,1.92) | 1.32(0.95,1.92) | 1.41(1.00,2.02) | 1.44(1.02,2.07) | <0.001 |
| Cholesterol,mmol/L | 5.07±1.13 | 5.16±1.14 | 5.13±1.12 | 5.12±1.13 | 5.01±1.13 | 4.95±1.13 | <0.001 |
| HDL,mmol/L | 1.32±0.34 | 1.33±0.37 | 1.34±0.34 | 1.33±0.34 | 1.30±0.33 | 1.28±0.34 | <0.001 |
| LDL,mmol/L | 2.98±0.90 | 3.03±0.91 | 3.04±0.89 | 3.02±0.89 | 2.94±0.90 | 2.88±0.89 | <0.001 |
| GGT,mmol/L | 21.0(15.0,32.0) | 22.0(16.0,33.0) | 21.0(15.0,32.0) | 21.0(15.0,32.0) | 21.0(15.0,33.0) | 21.0(15.0,33.0) | 0.4 |
| AST,mmol/L | 20.0(17.0,25.0) | 20.0(17.0,25.0) | 20.0(17.0,25.0) | 20.0(17.0,25.0) | 20.0(17.0,25.0) | 20.0(17.0,25.0) | 0.921 |
| ALT,mmol/L | 15.0(11.0,21.0) | 15.0(11.0,21.0) | 15.0(11.0,21.0) | 15.0(11.0,21.0) | 15.0(11.0,21.0) | 15.0(11.0,21.0) | 0.234 |
| FBG,mmol/L | 5.55(5.12,6.19) | 5.58(5.14,6.30) | 5.56(5.12,6.17) | 5.53(5.12,6.11) | 5.55(5.12,6.20) | 5.60(5.14,6.36) | <0.001 |
| PBG,mmol/L | 7.40(6.01,9.73) | 7.54(6.10,10.10) | 7.35(5.91,9.62) | 7.26(5.97,9.47) | 7.48(6.08,9.82) | 7.79(6.19,10.56) | <0.001 |
| Daytimenapping,% | 33.7(11393) | 10.6(69) | 12.5(358) | 18.9(3117) | 48.4(4307) | 71.5(3542) | <0.001 |
| Microalbuminuria,% | 13.3(4512) | 14.9(554) | 12.6(360) | 11.6(1912) | 14.6(1303) | 17.0(840) | <0.001 |
| High UACR level,% | 24.7(8357) | 26.7(174) | 26.0(741) | 23.8(3920) | 24.6(2189) | 26.9(1333) | <0.001 |
| CKD,% | 2.5(832) | 2.6(17) | 2.0(56) | 2.1(354) | 2.5(226) | 3.6(179) | <0.001 |
| Hyperfiltration,% | 2.5(860) | 2.8(18) | 2.3(65) | 2.2(363) | 2.8(250) | 3.3(164) | <0.001 |
| UACR,mg/g | 9.4(5.4,18.9) | 9.3(5.4,19.0) | 8.9(5.3,16.9) | 8.7(5.1,17.0) | 10.5(5.8,20.9) | 11.5(6.3,22.8) | <0.001 |
| eGFR(ml/(min*1.73m2) | 93.9±19.1 | 94.4±19.7 | 94.3±18.6 | 94.2±18.4 | 93.9±19.5 | 92.9±20.6 | <0.001 |

Chi-squared tests for discrete variables and one-way ANOVA for continuous variables
All continuous variables with normal distribution are presented as the mean values and standard deviation (SD). All continuous variables with skewness distribution are presented as media and 25,75 percentile. All enumeration data presented as proportion.
Microalbuminuria was defined as UACR>=30 mg/g. CKD was defined as eGFR<60 ml/min and hyperfiltration was defined as eGFR>=135 ml/min. According to the quartile division, among which the subjects belonged, the UACR data were divided into the under 25% group, the 25%-50% group, the 50%-75% group, and the over 75% group. Higher UACR level was defined as subjects belonged to the over 75% group.

Table 2 summarizes the participants' characteristics by daytime napping. Napping habit was reported by 7,001 of 11,198 men (62.5%) and 15,456 of 22,652 women (68.2%). Those who took naps seem to more likely to be women, which is converse to previous study^27^. Individuals who took naps were more likely to be regular drinkers, smokers and to have less physic exercise, naturally, they also have higher BMI and TG values and were more likely to have metabolic disease like obesity, hypertension, especially diabetes. Interestingly, napping seems to be a protective factor for hypercholesterolemia for the negative association between nap duration and CHOL, HDL. This result is consistent to the relationship between total sleep duration and CHOL, HDL. Nap habit also contributes to the kidney function decline. Participants took naps had higher prevalence of microalbuminuria, CKD and hyperfiltration, and their UACR values were significant higher.

**Table 2.**
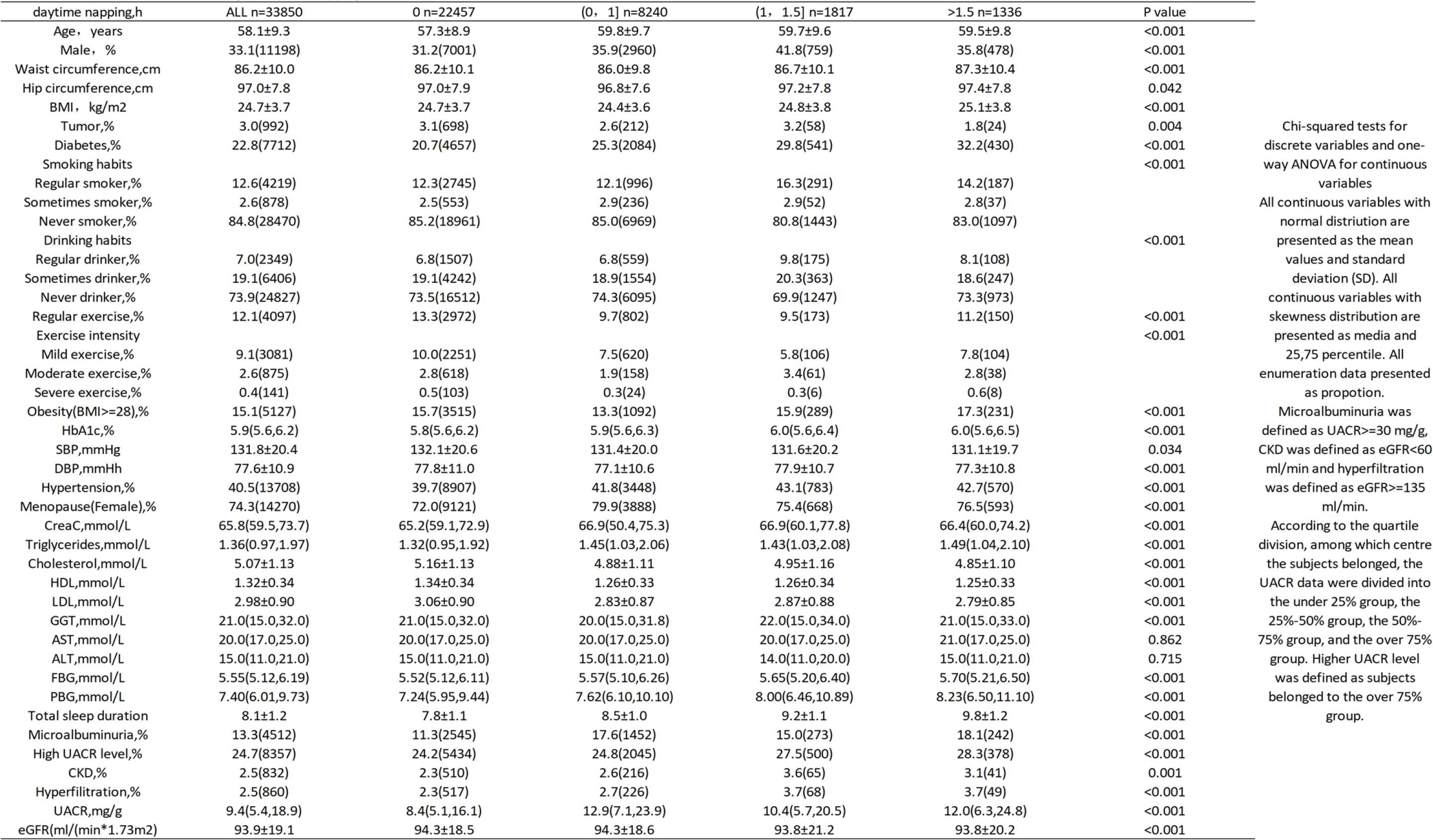
Baseline information according to daytime napping.

We further focus on the association between sleep duration and several health outcomes, including microalbuminuria, high UACR level, CKD, hyperfiltration, hypertension and diabetes. Multivariate logistic regressions were carried out before and after adjustment for possible confounding variables, such as age, sex, BMI, drinking and smoking habits, physic exercise, TG, CHOL, FBG, PBG, HbA1c, SBP, DBP, HDL, LDL, WC, HC. Results are shown at Table 3. Participants reporting short or long sleep duration had significant risks of microalbuminuria, high UACR level, hypertension and diabetes before adjustment in model 1, suggesting U-shaped relationship between sleep duration and health outcomes (Figure 1A). After adjusting for age, sex and BMI in model 2, the significant difference in hypertension outcome disappeared, suggesting differences in age, sex, and BMI between the sleep duration categories resulted in the higher risk for hypertension, and the relationship between short sleep duration and diabetes outcome had no statistical significance. After all confounding variables adjustment, only following several data were statistically significant. Compared with reference, fully adjusted ORs of >9 h/day for microalbuminuria and diabetes were 1.317 (1.200-1.446, p<0.001) and 1.288 (1.191-1.393, p<0.001); OR of (8, 9]h/day for microalbuminuria was 1.215 (1.123-1.315, p<0.001); ORs of <6h/day and [6, 7)h/day for high UACR level were 1.207 (1.045-1.392, p=0.010) and 1.126 (1.048-1.212, p=0.001). In addition, the OR of <6h/day for microalbuminuria was 1.218 (0.967-1.534, p=0.094), which was close to statistically significant. Therefore, these confounders had a strong interaction on the relationship between sleep duration and urinary protein, but the U-shaped trend relationship between sleep duration and urinary protein existed independent of these confounders (Figure 1C).

**Table 3.** Risk for health outcomes according to total sleep duration and daytime napping.

|  | microalbuminuria |  | higher UACR level |  | CKD |  | hyperfiltration |  | hypertension |  | diabetes |  |
| --- | --- | --- | --- | --- | --- | --- | --- | --- | --- | --- | --- | --- |
|  | OR | P value | OR | P value | OR | P value | OR | P value | OR | P value | OR | P value |
| Model 1 |  |  |  |  |  |  |  |  |  |  |  |  |
| Total sleep duration,h |  |  |  |  |  |  |  |  |  |  |  |  |
| (0, 6) | 1.335(1.070, 1.665) | 0.010 | 1.262(1.097, 1.452) | 0.001 | 1.222(0.747, 2.001) | 0.425 | 1.263(0.782, 2.042) | 0.339 | 1.183(1.010, 1.385) | 0.037 | 1.237(1.031, 1.484) | 0.022 |
| [6--7) | 1.101(0.976, 1.241) | 0.118 | 1.147(1.068, 1.231) | 0.000 | 0.912(0.686, 1.213) | 0.528 | 1.036(0.793, 1.352) | 0.798 | 1.006(0.928, 1.091) | 0.883 | 1.049(0.953, 1.155) | 0.329 |
| [7--8] | 1 |  | 1 |  | 1 |  | 1 |  | 1 |  | 1 |  |
| (8--9] | 1.308(1.212, 1.410) | 0.000 | 1.025(0.978, 1.074) | 0.296 | 1.188(1.003, 1.406) | 0.046 | 1.284(1.091, 1.511) | 0.003 | 1.059(1.004, 1.116) | 0.033 | 1.149(1.081, 1.223) | 0.000 |
| >9 | 1.558(1.427, 1.702) | 0.000 | 1.108(1.047, 1.174) | 0.000 | 1.710(1.425, 2.053) | 0.000 | 1.522(1.262, 1.836) | 0.000 | 1.149(1.078, 1.226) | 0.000 | 1.464(1.361, 1.574) | 0.000 |
| Daytime napping,h |  |  |  |  |  |  |  |  |  |  |  |  |
| 0 | 1 |  | 1 |  | 1 |  | 1 |  | 1 |  | 1 |  |
| (0, 1] | 1.674(1.560, 1.795) | 0.000 | 1.019(0.974, 1.067) | 0.398 | 1.158(0.986, 1.361) | 0.074 | 1.197(1.021, 1.402) | 0.026 | 1.095(1.040, 1.152) | 0.001 | 1.294(1.220, 1.373) | 0.000 |
| (1, 1.5] | 1.383(1.209, 1.584) | 0.000 | 1.120(1.028, 1.220) | 0.009 | 1.597(1.228, 2.076) | 0.000 | 1.650(1.275, 2.134) | 0.000 | 1.152(1.046, 1.269) | 0.004 | 1.621(1.458, 1.801) | 0.000 |
| >1.5 | 1.731(1.497, 2.001) | 0.000 | 1.113(1.008, 1.229) | 0.034 | 1.362(0.986, 1.882) | 0.061 | 1.616(1.199, 2.177) | 0.002 | 1.132(1.012, 1.266) | 0.030 | 1.814(1.610, 2.044) | 0.000 |
| Model 2 |  |  |  |  |  |  |  |  |  |  |  |  |
| Total sleep duration,h |  |  |  |  |  |  |  |  |  |  |  |  |
| (0, 6) | 1.251(1.000, 1.565) | 0.050 | 1.212(1.051, 1.395) | 0.008 | 1.065(0.645, 1.759) | 0.806 | 1.346(0.829, 2.184) | 0.229 | 1.058(0.892, 1.255) | 0.517 | 1.159(0.961, 1.398) | 0.122 |
| [6--7) | 1.083(0.959, 1.223) | 0.198 | 1.129(1.050, 1.212) | 0.001 | 0.887(0.665, 1.184) | 0.416 | 1.043(0.797, 1.365) | 0.758 | 0.961(0.880, 1.049) | 0.369 | 1.029(0.932, 1.136) | 0.570 |
| [7--8] | 1 |  | 1 |  | 1 |  | 1 |  | 1 |  | 1 |  |
| (8--9] | 1.264(1.171, 1.365) | 0.000 | 1.002(0.957, 1.050) | 0.927 | 1.031(0.868, 1.224) | 0.731 | 1.324(1.123, 1.561) | 0.001 | 1.000(0.944, 1.058) | 0.987 | 1.104(1.036, 1.177) | 0.002 |
| >9 | 1.442(1.318, 1.578) | 0.000 | 1.064(1.004, 1.126) | 0.035 | 1.273(1.054, 1.538) | 0.012 | 1.636(1.354, 1.977) | 0.000 | 1.005(0.937, 1.079) | 0.881 | 1.341(1.244, 1.446) | 0.000 |
| Daytime napping,h |  |  |  |  |  |  |  |  |  |  |  |  |
| 0 | 1 |  | 1 |  | 1 |  | 1 |  | 1 |  | 1 |  |
| (0, 1] | 1.561(1.453, 1.677) | 0.000 | 0.973(0.930, 1.019) | 0.244 | 0.887(0.752, 1.047) | 0.157 | 1.304(1.110, 1.530) | 0.001 | 0.948(0.896, 1.003) | 0.064 | 1.180(1.110, 1.255) | 0.000 |
| (1, 1.5] | 1.307(1.140, 1.499) | 0.000 | 1.112(1.020, 1.212) | 0.016 | 1.295(0.990, 1.693) | 0.059 | 1.731(1.333, 2.247) | 0.000 | 0.954(0.858, 1.060) | 0.379 | 1.427(1.279, 1.592) | 0.000 |
| >1.5 | 1.595(1.376, 1.849) | 0.000 | 1.053(0.954, 1.164) | 0.305 | 1.055(0.759, 1.468) | 0.749 | 1.761(1.302, 2.382) | 0.000 | 0.911(0.806, 1.029) | 0.132 | 1.619(1.431, 1.831) | 0.000 |
| Model 3 |  |  |  |  |  |  |  |  |  |  |  |  |
| Total sleep duration,h |  |  |  |  |  |  |  |  |  |  |  |  |
| (0, 6) | 1.218(0.967, 1.534) | 0.094 | 1.207(1.045, 1.392) | 0.010 | 1.100(0.663, 1.824) | 0.713 | 1.314(0.763, 2.261) | 0.325 | 1.014(0.851, 1.210) | 0.873 | 1.171(0.965, 1.421) | 0.110 |
| [6--7) | 1.088(0.960, 1.233) | 0.188 | 1.126(1.048, 1.212) | 0.001 | 0.934(0.699, 1.247) | 0.642 | 1.064(0.789, 1.435) | 0.683 | 0.950(0.868, 1.040) | 0.270 | 1.042(0.940, 1.154) | 0.436 |
| [7--8] | 1 |  | 1 |  | 1 |  | 1 |  | 1 |  | 1 |  |
| (8--9] | 1.215(1.123, 1.315) | 0.000 | 0.990(0.945, 1.039) | 0.688 | 1.030(0.866, 1.226) | 0.736 | 1.191(0.987, 1.438) | 0.068 | 0.974(0.918, 1.033) | 0.376 | 1.065(0.997, 1.138) | 0.060 |
| >9 | 1.317(1.200, 1.446) | 0.000 | 1.030(0.971, 1.092) | 0.322 | 1.216(1.002, 1.477) | 0.047 | 1.395(1.121, 1.736) | 0.003 | 0.974(0.906, 1.048) | 0.480 | 1.288(1.191, 1.393) | 0.000 |
| Daytime napping,h |  |  |  |  |  |  |  |  |  |  |  |  |
| 0 | 1 |  |  |  |  |  |  |  |  |  |  |  |
| (0, 1] | 1.477(1.370, 1.591) | 0.000 | 0.973(0.930, 1.020) | 0.267 | 0.847(0.715, 1.003) | 0.054 | 0.867(0.721, 1.041) | 0.126 | 0.966(0.911, 1.024) | 0.241 | 1.150(1.079, 1.226) | 0.000 |
| (1, 1.5] | 1.217(1.056, 1.403) | 0.007 | 1.079(0.988, 1.179) | 0.091 | 1.229(0.933, 1.619) | 0.143 | 1.265(0.933, 1.716) | 0.130 | 0.954(0.856, 1.064) | 0.401 | 1.434(1.279, 1.608) | 0.000 |
| >1.5 | 1.447(1.242, 1.687) | 0.000 | 1.035(0.934, 1.145) | 0.517 | 0.976(0.696, 1.367) | 0.886 | 1.188(0.837, 1.687) | 0.336 | 0.890(0.785, 1.009) | 0.890 | 1.623(1.427, 1.845) | 0.000 |

Model 1 was unadjusted. Model 2 was adjusted for age, sex and BMI. Model 3 was adjusted for covariates in model 2 plus Hip and waist circumference, drinking and smoking habits, physic exercise, triglycerides, cholesterol, HDL, LDL, GGT, blood glucose (except for diabetes) and blood pressure (except for hypertension).

**Figure 1.**
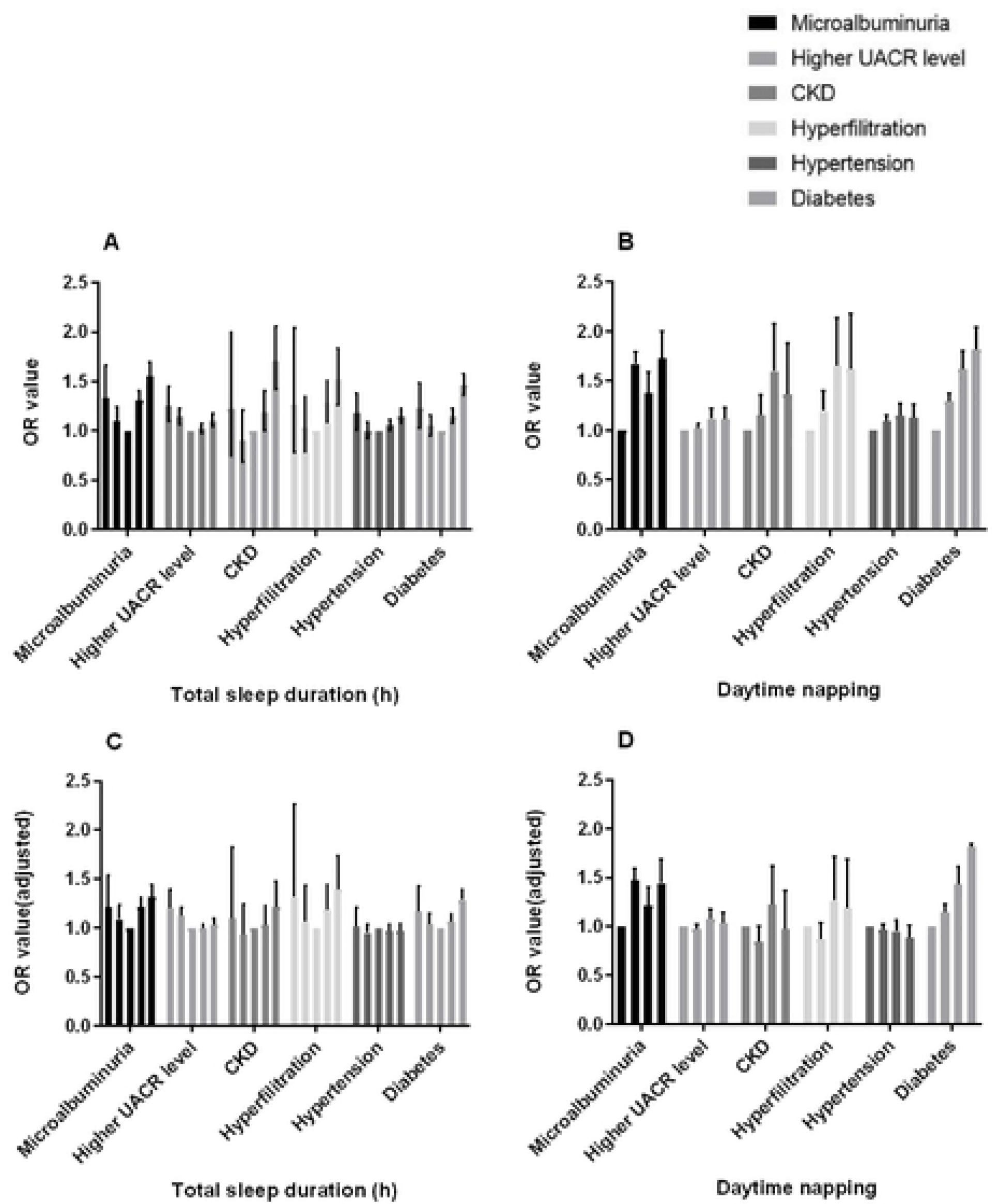
Association between sleep and risks for multiple health outcomes. (A&C) U-shaped association between total sleep duration and risks for multiple health outcomes. (B&D) Positive association between daytime napping and risks for multiple health outcomes. The bars in each group of data in the figure represent the total sleep duration of <6h, [6, 7)h, [7, 8]h, (8, 9]h and >9h, or represent the daytime napping of 0h, (0, 1]h, (1, 1.5]h and >1.5h successively. 1C and 1D were adjusted for age, sex, BMI, hip and waist circumference, drinking and smoking habits, physic exercise, triglycerides, cholesterol, HDL, LDL, GGT, blood glucose (except for diabetes) and blood pressure (except for hypertension).

According to the daytime napping categories, we found daytime napping duration is a risk factor for microalbuminuria and diabetes independent of confounders. Compared to subjects did not nap, the ORs of (0,1]h/day, (1,1.5]h/day and >1.5h/day for microalbuminuria were 1.477 (1.370-1.591, p<0.001), 1.217 (1.056, 1.403, p=0.007) and 1.447 (1.242, 1.687, p<0.001), for diabetes were 1.150(1.079, 1.226, p<0.001), 1.434(1.279, 1.608, p<0.001) and 1.623(1.427, 1.845, p<0.001) after fully adjustment (Figure 1B, 1D).

To further investigate relationship between total or daytime sleep duration and eGFR, CKD and hyperfiltration, we divided participants into eGFR>=90 ml/min group and eGFR<90 ml/min group. As we all known, in the progression of CKD, healthy individuals tended to have glomerular hyperfiltration initially, followed by increased risk for renal injury, leading to decrease in filtration rate and accelerating development of CKD. Thus we speculate eGFR level should be positively associated with inappropriate sleep duration among subjects with normal eGFR, while negatively associated with inappropriate sleep duration among subjects with lower eGFR. The results confirmed our conjecture. eGFR value was U-shaped associated with sleep duration among normal eGFR group (Figure 2A), and N-shaped associated with sleep duration among lower eGFR group (Figure 2B), suggesting both short and long sleep duration contributes to the progression of kidney function decline, which was consistent to the urinary protein-sleep duration relationship. Multivariate logistic regressions were carried out to examine the results, as it shown at Table 4, after fully adjustment, OR of >9h/day sleep for hyperfiltration was 1.400 (1.123-1.745, p=0.003) among eGFR>=90 group. The U-shaped trend relationships between sleep duration and hyperfiltration among normal eGFR subjects or CKD among lower eGFR subjects exist (Figure 2C), though the statistical significance not ideal. It could be explained by few of our participants had eGFR<90 ml/min.

**Table 4.**
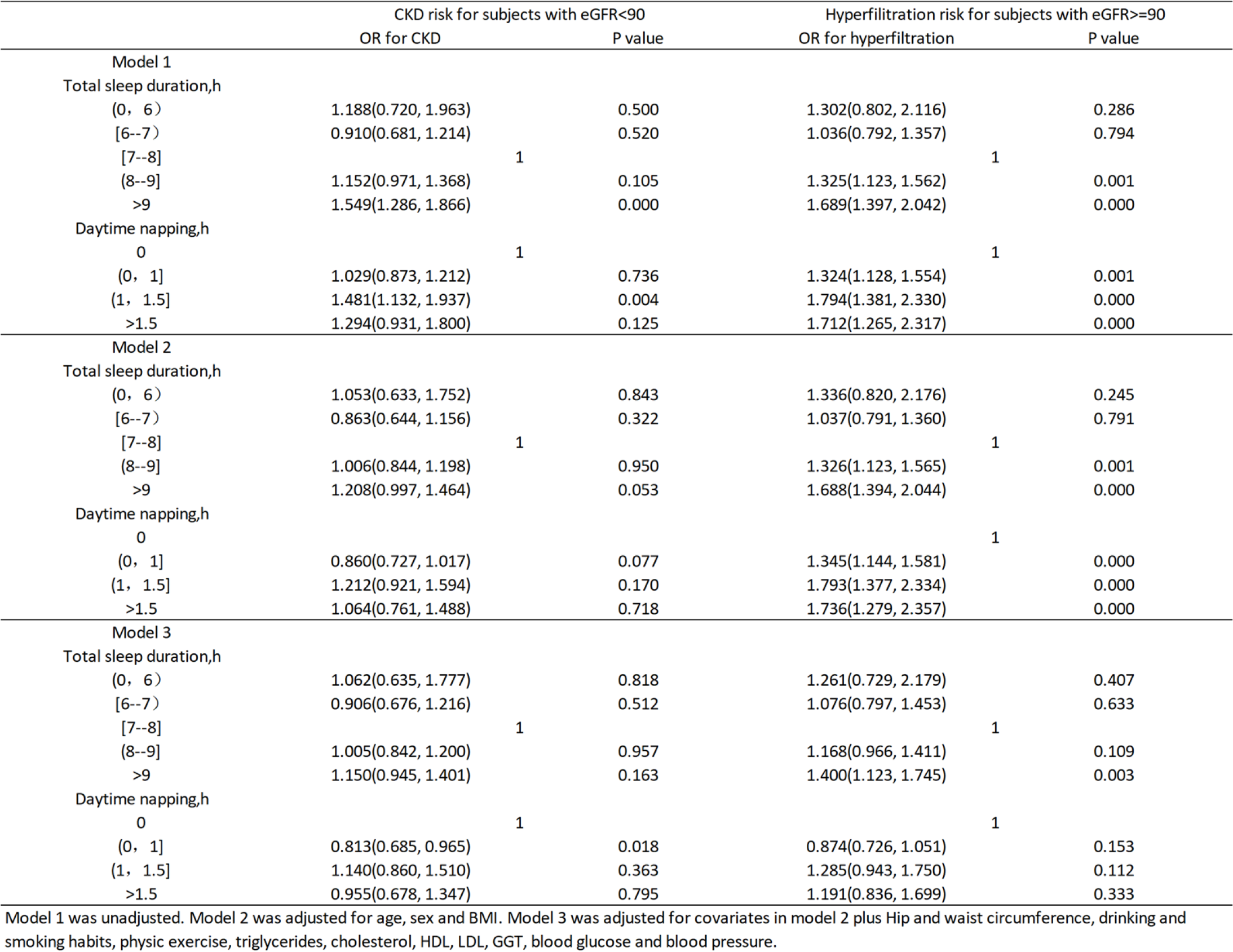
Relationships between total sleep duration, daytime napping and eGFR.

**Figure 2.**
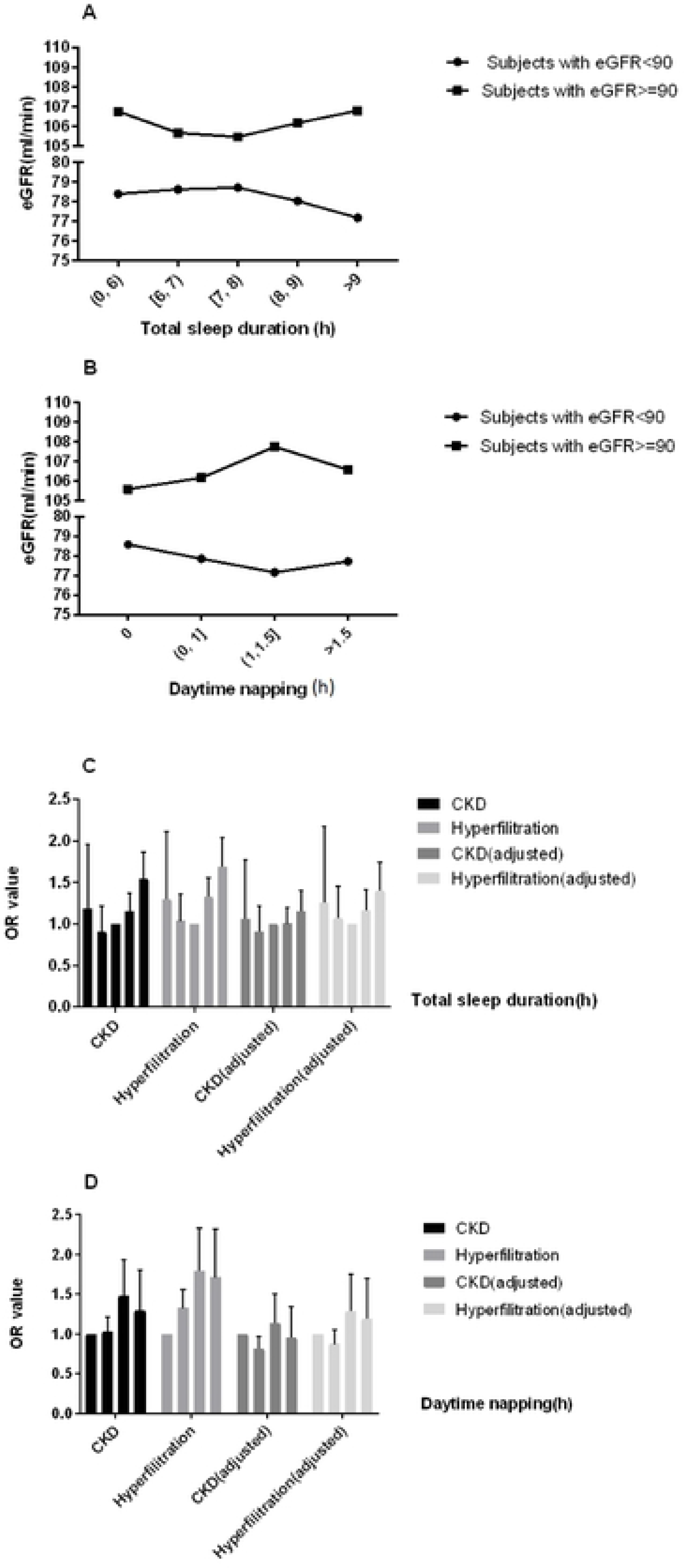
Association between sleep and eGFR. (A) Association between mean eGFR values and total sleep duration among subjects with <90 ml/min and >=90ml/min eGFR. (B) Association between mean eGFR values and daytime napping among subjects with <90 ml/min and >=90ml/min eGFR. (C&D) Risks for hyperfiltration among subjects with >=90ml/min eGFR, and risks for CKD among subjects with <90ml/min eGFR, according to total sleep duration or daytime napping. Adjusted covariates include age, sex, BMI, hip and waist circumference, drinking and smoking habits, physic exercise, triglycerides, cholesterol, HDL, LDL, GGT, blood glucose and blood pressure. CKD was defined as eGFR<60 ml/min and hyperfiltration was defined as eGFR>=135 ml/min.

We noticed that daytime napping has a positive effect on the risk for hyperfiltration among eGFR>=90 ml/min group in model 1 and model 2 (Table 4, Figure 2D). However, after fully adjustment, the statistical significance disappear, suggesting daytime napping affects the occurrence of hyperfiltration by influencing confounding factors such as blood glucose, BMI and blood lipid. It shall be supported by our previous finding that napping significantly increased the risk of diabetes (Table 3).

Actually, total sleep duration and daytime napping duration have an interaction on the risk of health outcomes as well. Subjects who sleep for a short time usually do not nap while those who sleep for a long time often have napping habit. Therefore, we conducted a joint analysis to investigate the interaction. We divided participants into 20 subgroups according to their total sleep duration and daytime napping. Following five groups were excluded from the reanalysis because there were fewer than 100 people: total sleep duration<6h&daytime napping between 0 and 1h; total sleep duration<6h&daytime napping between 1 and 1.5h; total sleep duration<6h&daytime napping>1.5h; total sleep duration between 6 and 7h&daytime napping between 1 and 1.5h; total sleep duration between 6 and 7h&daytime napping>1.5h. A multivariate logistic regression after fully adjustment was carried out and result is shown in Figure 3, several results were notable. First, subjects with [7, 8]h total sleep duration and did not had a napping habit have the lowest risk for microalbuminuria. Second, The U-shaped relationship of risk for microalbuminuria and sleep duration was significant in the non-napping group, and the ORs of almost all these groups reached the statistically significant level (P=0.017, 0.081, 0.003, 0.000 respectively). This result not only indicated the U-shaped curve relationship more convincing, but also explained the reason why the statistical significance of short sleep duration groups was poor in our previous logistic regression analysis. It was because the subjects with short total sleep duration often do not have napping habit, and napping is also a risk factor for microalbuminuria, which prevent them from having a significant higher risk for microalbuminuria compared to the reference. Third, subjects with [6, 7)h total sleep duration and (0, 1]h daytime napping had the highest risk for microalbuminuria (OR=1.959, P<0.001). This was the group with the shortest total sleep duration and the longest napping that we included in the analysis (groups with shorter total sleep duration and longer napping were excluded for small sample size, and the P values of these groups were poor.), suggesting a lack of nighttime sleep can be dangerous for people who are already short on total sleep duration, and daytime napping is not enough to make up for the loss of sleep at night, which is consistent to conjecture of Miao et al^28^. Fourth, subjects with the longest total sleep duration and longest napping duration had the second high risk for albuminuria (OR=1.625, P<0.001), suggesting longer napping duration will further aggravate the risk of extremely long total sleep duration for microalbuminuria.

**Figure 3.**
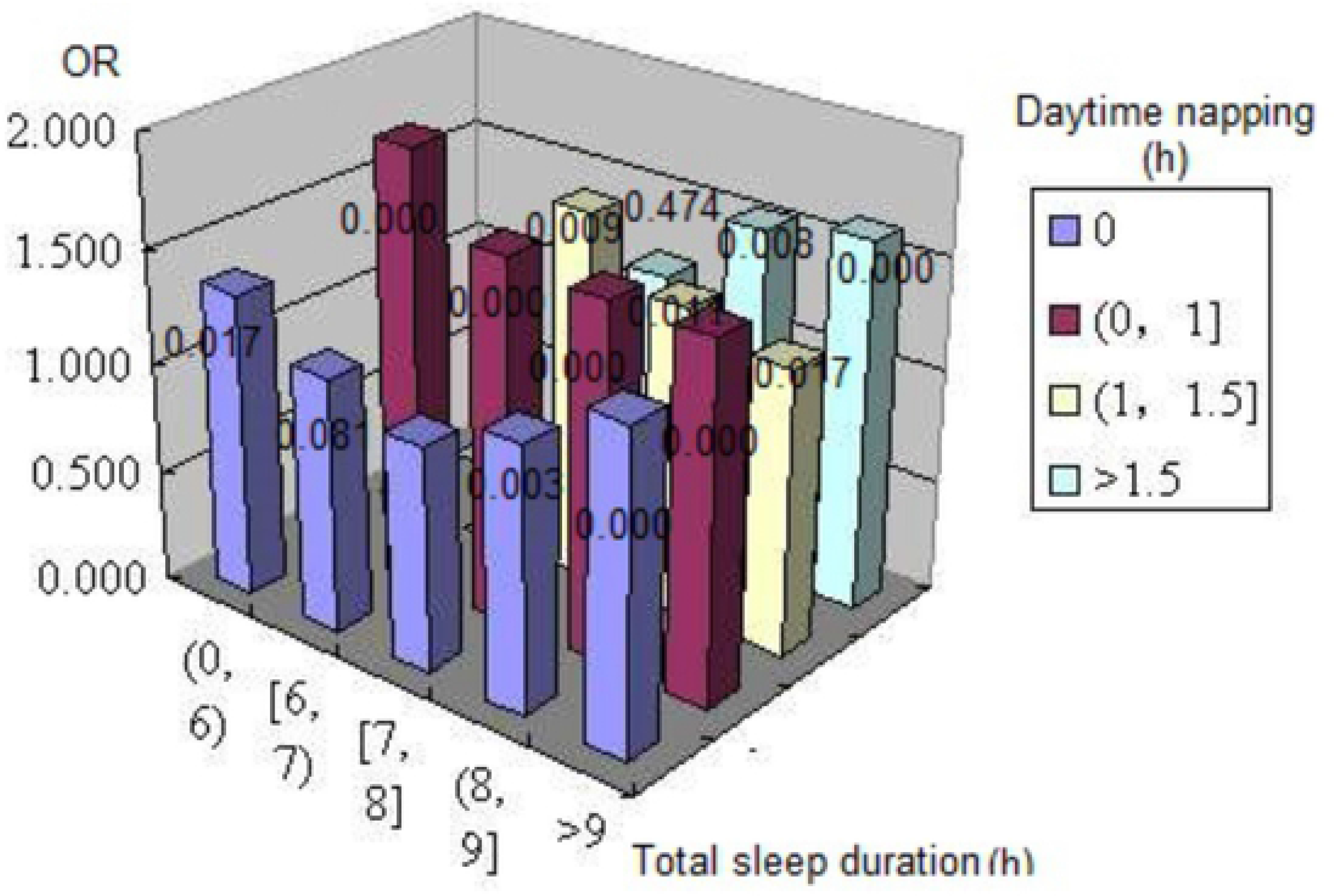
Joint analysis on total sleep duration and daytime napping in relation to the risk for microalbuminuria. Multivariate ORs for microalbuminuria were adjusted for age, sex, BMI, hip and waist circumference, drinking and smoking habits, physic exercise, triglycerides, cholesterol, HDL, LDL, GGT, blood glucose and blood pressure. Asterisk denotes result statistically different from [7, 8] hours of sleep duration per day with out daytime napping.

At last, we investigated the interactions of several confounders, which was commonly considered to be the main risk factor for microalbuminuria, including blood pressure, blood glucose and BMI, the results are shown at Table 5. We found long sleep duration is a statistically significant independent risk factor for microalbuminuria among subjects with >100 mmHg SBP and <90 mmHg DBP, but not among subjects with <=100 mmHg SBP or >=90 mmHg DBP. This may be due to the fact that blood pressure is the main factor affecting urinary protein, and the effect of sleep duration on urinary protein is insignificant for subjects with extremely poor or excellent blood pressure control. According to the results of different categories of glucose metabolism and BMI, both blood glucose and BMI had an interaction on the sleep-albuminuria association. For subjects with abnormal glucose metabolism or obesity problem, poor sleep habits were even worse for their health. This could be explained by our previous finding that extremely long sleep duration shall aggravate their obesity or diabetes, followed by worse effects that long sleep duration bring to their health, which become a vicious cycle.

**Table 5.** Risk for microalbuminuria according to total sleep duration among diverse subgroup.

|  | Total sleep duration, h |  |  |  |  |
| --- | --- | --- | --- | --- | --- |
|  | (0, 6) | [6--7) | [7--8] | (8--9] | >9 |
| **SBP(mmHg)** |  |  |  |  |  |
| ≤100 | 1.335(0.666) | 0.844(0.700) | 1 | 1.198(0.499) | 0.912(0.787) |
| 100-140 | 1.333(0.069) | 1.089(0.327) | 1 | 1.212(0.000) | 1.277(0.000) |
| ≥140 | 1.043(0.815) | 1.098(0.333) | 1 | 1.184(0.006) | 1.352(0.000) |
| **DBP(mmHg)** |  |  |  |  |  |
| <90 | 1.278(0.054) | 1.114(0.125) | 1 | 1.217(0.000) | 1.340(0.000) |
| ≥90 | 0.883(0.675) | 0.949(0.724) | 1 | 1.135(0.180) | 1.063(0.848) |
| **glycometabolism** |  |  |  |  |  |
| NGT | 1.287(0.179) | 1.088(0.402) | 1 | 1.180(0.009) | 1.189(0.028) |
| IFG/IGT | 1.192(0.425) | 1.023(0.850) | 1 | 1.286(0.001) | 1.266(0.008) |
| DM | 1.319(0.176) | 1.159(0.196) | 1 | 1.216(0.008) | 1.552(0.000) |
| **BMI(kg/m<sup>2</sup>)** |  |  |  |  |  |
| <24 | 0.846(0.436) | 0.999(0.995) | 1 | 1.199(0.003) | 1.296(0.000) |
| 24-27 | 1.227(0.333) | 1.211(0.085) | 1 | 1.199(0.012) | 1.313(0.001) |
| 27-30 | 1.915(0.008) | 1.161(0.298) | 1 | 1.317(0.004) | 1.311(0.018) |
| ≥30 | 1.337(0.370) | 0.946(0.789) | 1 | 1.172(0.255) | 1.497(0.015) |

The model was adjusted for covariates in model 2 plus Hip and waist circumference, drinking and smoking habits, physic exercise, triglycerides, cholesterol, HDL, LDL, GGT, blood glucose and blood pressure. The data are presented as OR(P values).

## Discussion

In this study, we identified associations between total sleep duration, daytime napping, and the incidence of microalbuminuria, high UACR level, CKD and hyperfiltration in a community middle-aged Chinese population. We found that total sleep duration was U-shaped associated with the incidence of microalbuminuria and the progression of kidney function decline, and daytime napping was associated with the incidence of microalbuminuria positively. It was speculated extreme long sleep duration could significantly aggravate the process of kidney damage, lead to hyperfiltration first and then eGFR decline. Further joint analysis showed the importance for the kidney function outcomes that people with short sleep duration should ensure enough nighttime sleep and people with long sleep duration should limit the daytime napping.

Several studies had investigated the association between sleep and kidney health outcome. Both short and long sleep duration were reported to be associated with decreased eGFR and the progression to ESRD among CKD^12, 21, 32, 33^ or hypertension^18^ population, and to be associated with increased eGFR, hyperfiltration, inadequate hydration and the prevalence of CKD among community-based general population^13, 19, 20, 26–28^. In our study, based on same general community population, we further demonstrated that inappropriate sleep duration had converse effects on eGFR in healthy or early-stage nephropathy population, which was consistent to previous studies. However, a US study of 4,238 participants from Nurses' Health Study (NHS) reported short sleep duration was prospectively associated with faster decline in kidney function among a healthy general population^11^. An explanation for that is NHS study was based on subjects whose jobs were nurse, which is often be high-intensity workers and shift workers. A Japanese study suggested that inappropriate sleep duration was more likely to affect kidney health and raise the risk of early stage kidney disease for shift workers^34^. Indeed, the participants of NHS study had a lower mean eGFR (88.3±25.0) than our participants (93.9±19.1), therefore, their results were more similar to those in the CKD population. A few studies selected albuminuria as a terminal outcome to evaluate how sleep affects kidney function. Both short and long sleep duration had been reported to be associated with UACR level among populations from Japan^35^, Korea^36^ and US^12^, and daytime napping had been reported to be positively associated with albuminuria in Japan^37^, however, this relationship is racial-specific^13^, whether it exists among Chinese population keeps unknown. This study confirmed the U-shaped relationship between sleep duration and albuminuria and the positive relationship between napping and albuminuria on the basis of the Chinese population. In addition, our samples were from 8 different regions in China, including coastal and inland regions, developed and underdeveloped regions, with a wide geographical span, which is an ideal representation of the Chinese population. Finally, we investigated the interaction between total sleep time and napping, total sleep time and blood pressure, blood glucose and BMI, and provided a reference for individuals with diverse specific conditions to control appropriate sleep duration.

The correlation between sleep and kidney function can be explained in the following ways: First, both short and long sleep duration shall result in systemic inflammation, which may account mainly for the association with increased UACR levels. Long sleep duration is associated with subclinical inflammation and increased arterial stiffness^38–41^, while sleep curtailment increases the proinflammatory cytokines^42^, high-sensitivity C-reactive protein^43^ and white blood cell^44^ levels, which reflects systemic inflammation, causes glomerular endothelial dysfunction and consequently leads to albuminuria^45, 46^. Second, changes in sympathetic nervous system influenced by higher or lower sleep duration may cause kidney function decline^47^. Sleep regulates the activity of hypothalamic pituitary adrenal (HPA) axis. Activity of HPA axis is reduced during sleep onset and early stages of sleep, while it is activated during latter stages of sleep, such as rapid eye movement (REM) stage^41, 48, 49^. Therefore, extreme short sleep may weaken the inhibitory effect of early stage sleep on the HPA axis, while extreme long sleep may enhance the activation effect of REM stage on the HPA axis, thus keeping the activity of the HPA axis at higher level, which is adverse to the metabolic health. Third, sleep deprivation in humans reduces plasma renin, angiotensin and aldosterone levels, which is associated with increased urinary excretion of sodium and potassium^50, 51^. In addition, the normal nocturnal dipping of blood pressure is attenuated. Related animal experiments had proved that sleep deprivation caused increased sympathetic nerve activity and reduced plasma angiotensin Ⅱ levels^52^. Fifth, regarding the association between napping and microalbuminuria, changes in the circadian rhythms was speculated to contribute to the effect of sleep on kidney function. Animal models were constructed by mutating the circadian regulatory gene casein kinase-1ε, related experiment showed animals’ heterozygote for the mutation exhibit phase-advanced and shortened circadian rhythms and were shown to develop albuminuria, renal tubular atrophy and cardiac dysfunction^53^. This is supported by our findings that daytime napping could not compensate for lack of sleep at night when total sleep duration was equal, and by results reported by Sasaki et al^34^. That short sleep duration was more likely to affect kidney health in shift workers. Fifth, different distributions of sleep-disordered breathing (SDB)^54–57^ and restless leg syndrome (RLS)^46, 47, 55^ in several sleep duration groups may also be a reason, according to previous reports.

The major strength of our study is large sample of the general population from three areas of China, which made our results representative and statistically significant, also provided the possibility for our subsequent subgroup analysis of interactions. REACTION study was an epidemiological investigation on tumors. In addition, detailed medical history and drug use history were collected during the questionnaire process. In this way, subjects with serious diseases and using ACEI/ARBs drugs could be excluded in the subsequent statistics to obtain more reliable results. However, current study also has several limitations. First, sleep duration was determined according to a self-reported questionnaire, as in many prior epidemiological studies and was not measured objectively; additionally, sleep quality cannot be evaluated, and prevalence of SDB or RLS could not acquire. Second, the cross-sectional study design prevents us from establishing a causal relationship between sleep and kidney function. Third, protein content intake the previous day and the interval between last meal and sleep were not available, which may affect UACR. Forth, UACR levels were determined by a single measurement, but detection methods of the 8 centres were different; hence, we set several outcomes, and the values of UACR were divided according to the quartile division in the centre to which the subject belonged, to estimate UACR level in logistic regression.

In summary, the goal of our study is demonstrating the relationship between sleep duration, daytime napping, UACR and eGFR in middle-aged apparently healthy Chinese population. We provided an explanation for the current epidemiological investigation into the controversial inconsistent results of the relationship between sleep duration and eGFR levels. Further cohort study should to be done to confirm our conclusions.

## Acknowledgments and Financial Disclosures

The authors thank Jia Li, Yingfei Zhu, Yang Yu, Tianyu Xie from Nankai University and Haibin Wang from Medical School of Chinese PLA for Statistical knowledge consultation. No potential conflicts of interest relevant to this article were reported.

Present study was supported by Chinese Society of Endocrinologoy, the Key Laboratory for Endocrine and Metabolic Diseases of Ministry of Health (1994DP131044), the National Key New Drug Creation and Manufacturing Program of Ministry of Science and Technology (2012ZX09303006-001), the National High Technology Research and Development Program of China (863 Program, 2011AA020107). National Science and Technology Major Project 288 (2011ZX09307-001-08).

